# Exploiting *Pseudomonas syringae* Type 3 secretion to study effector contribution to disease in spinach

**DOI:** 10.1101/2024.06.14.599008

**Authors:** Melanie Mendel, Xander C. L. Zuijdgeest, Femke van den Berg, Leroy van der Meer, Joyce Elberse, Petros Skiadas, Michael F Seidl, Guido Van den Ackerveken, Ronnie de Jonge

## Abstract

Intensive spinach cultivation creates favourable conditions for the emergence and rapid evolution of pathogens, leading to substantial economic losses. Research on host-pathogen interactions in leafy greens would benefit from advanced biotechnological tools, however absence of such tools in spinach hampers our understanding of spinach immunity. Here, we explored the potential of Type III Secretion System (T3SS)-mediated effector delivery to study pathogen effector activity in spinach. We identified the *Pseudomonas syringae* pv*. tomato* DC3000 (DC3000) polymutant D36E, which lacks 36 known T3SS effectors (T3Es), as a promising T3SS-dependent effector delivery system for spinach. Unlike DC3000, which causes necrotic symptoms on spinach and reaches high bacterial titres, D36E did not proliferate and caused no visible symptoms. Using D36E, we screened 28 DC3000 T3Es in spinach, assessing symptom development, bacterial proliferation, and reactive oxygen species (ROS) bursts as a proxy for early immune responses. AvrE1 and HopM1 emerged as key determinants of DC3000-like infection, inducing water-soaked lesions, while HopAD1 strongly suppressed ROS production. Our findings establish the D36E-based effector delivery system as a powerful tool for high-throughput effector studies in spinach. It bridges the gap between genomics-based effector predictions and experimental validation, paving the way for knowledge-driven resistance breeding in non-model crops like spinach.

## Introduction

As consumers increasingly opt for healthier dietary alternatives, spinach has seen a remarkable rise in its economic significance (Kandel et al., 2019). To meet the increasing demand, growers cultivate spinach at high density, providing favourable conditions for the outbreak and rapid evolution of pathogens. The oomycete *Peronospora effusa* (*P. effusa*) poses a major threat to global spinach production. It infects spinach leaves, and causes downy mildew disease, rendering the crop unsellable and resulting in substantial economic losses (Ribera et al., 2020). Other agronomically relevant spinach pathogens include the oomycetes *Albugo occidentalis, Pythium spp.*, the fungus *Colletotrichum dematium,* and viruses like the cucumber mosaic virus (Kandel et al., 2019; Ribera et al., 2020). To mitigate the impact of these pathogens on spinach production, targeted improvement of crop resistance is crucial. However, unlike other leafy greens such as lettuce, spinach lacks biotechnological tools to study its immunity, limiting our ability to identify and develop novel disease resistance traits efficiently.

Our understanding of the spinach immune system remains limited and is primarily based on comparative genome analysis and transcriptomics of disease resistance. For example, a comparison of *P. effusa*-susceptible and *P. effusa*-resistant spinach cultivars transcriptomes showed that defence-related genes like *PR-1* and diverse types of immune receptor genes, including the receptor-like kinase *FERONIA*, are upregulated in resistant spinach cultivars (Kandel et al., 2020; Clark et al., 2024). *FERONIA* is hypothesized to play a role in plant immunity, including the downstream production of reactive oxygen species (ROS), a key component of plant immunity (Cheung, 2024). Immune receptors such as the receptor-like kinases (RLKs) and nucleotide-binding leucine-rich repeat receptors (NLRs) are involved with a variety of immune responses including, programmed cell death also known as the hypersensitive response (HR), preventing the spread of pathogens (Jones and Dangl, 2006; Ngou et al., 2022). The spinach genome encodes several hundred NLRs, of which the role in pathogen detection is largely unknown (Xu et al., 2017; Hulse-Kemp et al., 2021).

Likewise, we know little about the molecular disease-related processes that are important during spinach-pathogen interactions, including the activity and targets of pathogen effectors. Only recently, genomic resources for spinach and its pathogens became available (Hulse-Kemp et al., 2021; Fletcher et al., 2022, Skiadas et al., 2024). Nevertheless, no effector has been thus far identified to be recognized by spinach (Cai et al., 2021). Identifying effectors and corresponding immune receptors would greatly benefit future research and breeding efforts for disease-resistant spinach but requires high-throughput methods to screen large number of putative effectors. To screen for effector recognition, typically, *Agrobacterium*-mediated transient expression assays are used (Lu et al., 2016; Pelgrom et al., 2019). Recent reports on improved *Agrobacterium*-mediated transient expression in spinach focused on fine-tuning of individual protein expression levels but were not designed to detect immunity- or pathogenicity-related phenotypes (Zhang et al., 2024). Alternatives approaches to deliver effector candidates into plants include biolistic methods, nanoparticles, chemical transformations, fusion peptides, and the bacterial Type 3 Secretion System (T3SS), however, to date, none of these have been explored in spinach for effector research (Rustgi et al., 2022). Here, we evaluated the potential of the T3SS as a method for studying pathogen effectors in spinach. The T3SS, a molecular syringe, is essential for the delivery of bacterial effectors into the host cell (O’Malley and Anderson 2021). T3SS mutants typically display reduced virulence, underscoring the importance of T3SS-delivered effectors (T3Es) in establishing disease (Yuan and He 1996). Using the effectorless *Pseudomonas syringae* pv. *tomato* DC3000 polymutant D36E, we here show that in the absence of T3Es, D36E survives in spinach leaves while causing no detectable symptoms. To verify if D36E can deliver effectors in spinach, we reintroduced 28 known DC3000 T3Es into D36E and evaluated theircontribution to symptom development, bacterial proliferation, and ROS burst alterations, an early marker of plant immunity. We demonstrate the effects of several of these effectors on these phenotypes in spinach, supporting previous evidence that these effectors target conserved components of plant immunity and physiology.￼

## Results

### Identification of *Pseudomonas* strain D36E as a candidate for T3SS-dependent effector delivery in spinach

To study effectors in spinach, we first sought to identify a T3SS-encoding bacterial strain that is suited for the delivery of effectors through the T3SS into spinach. As initial indicator of a bacterium’s suitability for T3SS-dependent effector delivery, we tested for the absence of visible changes at the infiltration site, and the bacterium’s ability to sustain a population when inoculated into spinach leaves (Figure 1). We tested multiple *Pseudomonas* strains that were previously employed to deliver effectors in other hosts, i.e.: 1) the Effector-to-Host-Analyser strain (EtHAn), a non-pathogenic *Pseudomonas fluorescence* (*Pf-*0) strain that had been genetically engineered to express a T3SS and has been successfully employed in multiple plants across several studies (Thomas et al., 2009; Krasileva et al., 2011; Upadhyaya et al., 2013); 2) the wildtype *Pseudomonas syringae* pv*. tomato* DC3000 (DC3000) strain, a natural pathogen of tomato and that has been used to study effectors in other plants (Buell et al., 2003; Fabro et al., 2011); 3) multiple DC3000 mutants with reduced effector arsenals, i.e., the DC3000^Δ*CEL*^(Δ*CEL*) mutant that lacks a conserved effector locus, the nearly effectorless DC3000^D28E^ (D28E) polymutant, and the effectorless polymutant DC3000^D36E^ (D36E) (Alfano et al., 2000; Cunnac et al., 2011; Wei et al., 2015); and 4) a *Pseudomonas syringae* pv*. phaseolicola* (*Pph*) strainthat is a natural pathogen of bean and has displayed promising results in preliminary work on spinach (Arnold et al., 2011). Lastly, we included a fourth DC3000 mutant, the T3SS mutant DC3000^Δ*hrcC*^ (Δ*hrcC*) that cannot assemble a functional T3SS and is therefore unable to deliver T3Es (Yuan and He, 1996).

**Figure 1:**
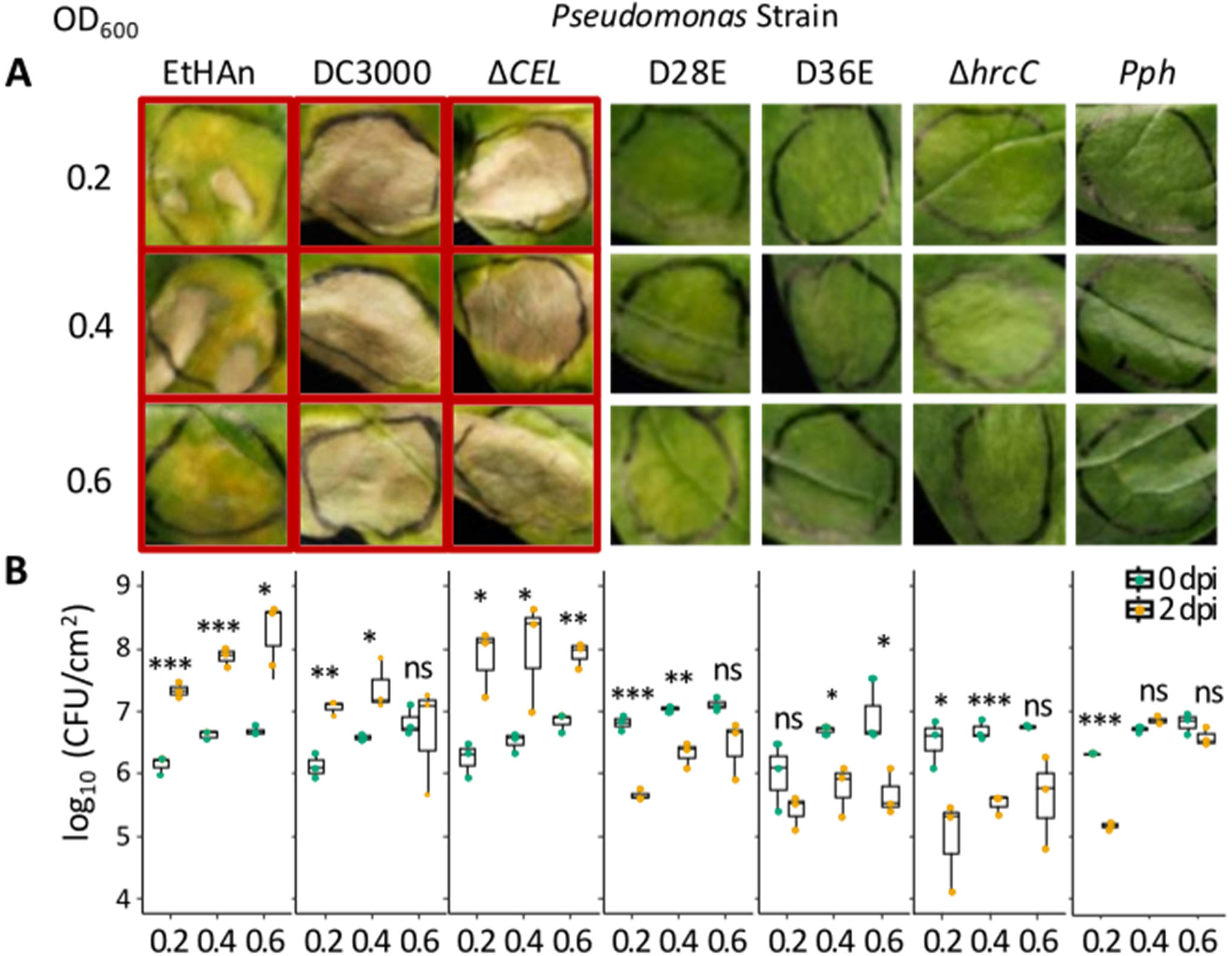
*Pseudomonas syringae* pv. *tomato* DC3000 polymutant D36E is a suitable candidate for effector screens in spinach. **A)** Representative images of spinach leaves 6–7 days post infiltration (dpi) with various *Pseudomonas* strains: the transgenic *Pseudomonas fluorescens* Pf-1 strain EtHAn (EtHAn), *Pseudomonas syringae* pv. *tomato* DC3000 (DC3000), the DC3000 mutant strain Δ*CEL* (Δ*CEL*), the DC3000 effector polymutants D28E (D28E) and D36E (D36E), the DC3000 T3SS-mutant Δ*hrcC* (Δ*hrcC*) and the bean pathogen *Pseudomonas syringae* pv. *phaseolicola* (*Pph*). Photographs were taken 67 days post inoculation (dpi). Infiltration sites outlined in red highlight severe chlorosis and necrosis, particularly in EtHAn-, DC3000-, and Δ*CEL*-infiltrated leaves. **B)** Bacterial proliferation quantified as log_10_ colony-forming units (CFU) per square centimetre (cm^2^) at 0 dpi (green; immediately after infiltration) and 2 dpi (yellow). Infiltrations in **(A)** and **(B)** were performed on the first true leaves of 2.5-3-week-old spinach plants at the indicated inoculum densities (0D_600_ = 0.2, 0.4, and 0.6). Statistical significance between CFU counts at 0 dpi and 2 dpi was determined using a two-tailed t-test (*** *p* < 0.001, ** *p* < 0.01, * *p* < 0.05; *n* = 3 infiltration sites per treatment and timepoint).

We syringe-infiltrated the first true leaves of 2.5-3-week-old spinach with *Pseudomonas* bacteria at high optical densities at 600 nm (OD_600_) equal to 0.2, 0.4 and 0.6. We then visually assessed infiltration sites between 6 days post inoculation (dpi) and 7 dpi, confirming that Δ*hrcC* and *Pph* did not cause visible symptoms at the infiltration site. EtHAn, DC3000, and Δ*CEL* consistently caused strong necrotic symptoms at all tested concentrations and D28E induced only mild chlorosis at the highest concentrations, while D36E caused no visible symptoms (Figure 1A). To estimate the corresponding bacterial populations in spinach, we next measured bacterial titres in spinach as colony-forming units (CFUs) at 0 dpi and 2 dpi (Figure 1B). The Δ*hrcC* and D36E strains exhibited 1.5- to 2-fold reductions in bacterial titres at 2 dpi compared to 0 dpi. *Pph*, like Δ*hrcC,* showed reduced bacterial titres between 0 dpi and 2 dpi at an OD_600_ of 0.2. However, at higher inoculum densities (OD_600_ of 0.4 and 0.6), *Pph* maintained bacterial titres at 2 dpi close to 0 dpi levels, suggesting that native *Pph* effectors may contribute to titre stabilization. EtHAn, DC3000 and Δ*CEL*, all of which exhibited necrotic symptoms, proliferated in spinach between 0 dpi and 2 dpi, increasing titres around 1.5-2 orders of magnitude. In contrast, D28E and D36E behaved like Δ*hrcC*, with titres 1-1.5 orders of magnitude lower at 2 dpi compared to 0 dpi. To assess whether D36E sustains a population in spinach after 2 dpi, we further monitored its titres in leaves over a period of five consecutive days (Figure S1). We observed that D36E titres stabilized after 1 dpi, indicating that D36E can persist in spinach for at least five days without eliciting visible plant responses (Figure S1). Since the effectorless D36E polymutant did not induce visible alterations yet encodes a functional T3SS to deliver effectors, we concluded that this strain is suitable for effector delivery into spinach.

### AvrE1 and HopM1 are key contributors to visible DC3000-like disease symptoms in spinach

Since infiltration of wildtype DC3000 induced necrotic symptoms in spinach, whereas D36E did not (Figure 1), we hypothesised that these symptoms result from the activity of specific DC3000 effectors. To test this, we assessed D36E’s suitability as an effector delivery system by individually reintroducing 28 DC3000 effectors into D36E, generating a library of D36E strains, each carrying a single effector. We then evaluated the impact of these effectors on symptom development at infiltration sites and bacterial titres in spinach.

To systematically describe and categorise visual symptoms in spinach, we used a semi-quantitative scoring matrix ranging from 0 (no symptoms, D36E-like) to 4 (strong necrosis, DC3000-like). While representative images illustrate these symptoms, considerable variation was observed (Figures 2, S2). Among the 28 tested single-effector D36E strains, 18 did not induce consistent symptoms beyond those of D36E. In contrast, 10 effectors triggered diverse visible symptoms. The clearest symptoms were seen with *AvrE1* and *HopM1,* which caused strong necrosis in over 80% of infiltration sites (symptom score >2), closely followed by *HopAM1*, which also produced symptoms similar to those caused by DC3000. *HopF2* and *HopN1* also resulted in a disease-score >1 in at least 65% of infiltration sites, though with variable frequency and severity. For example, *HopF2* predominantly induced chlorosis, with necrotic lesions occurring in fewer than half of the infiltration sites. Infiltration of D36E carrying *AvrPtoB*, *HopAD1*, *HopAO1*, *HopR1*, and *HopU1* significantly increased symptom frequency, though symptom scores >1 were inconsistent, appearing in only 27%-47% of infiltration sites.

**Figure 2:**
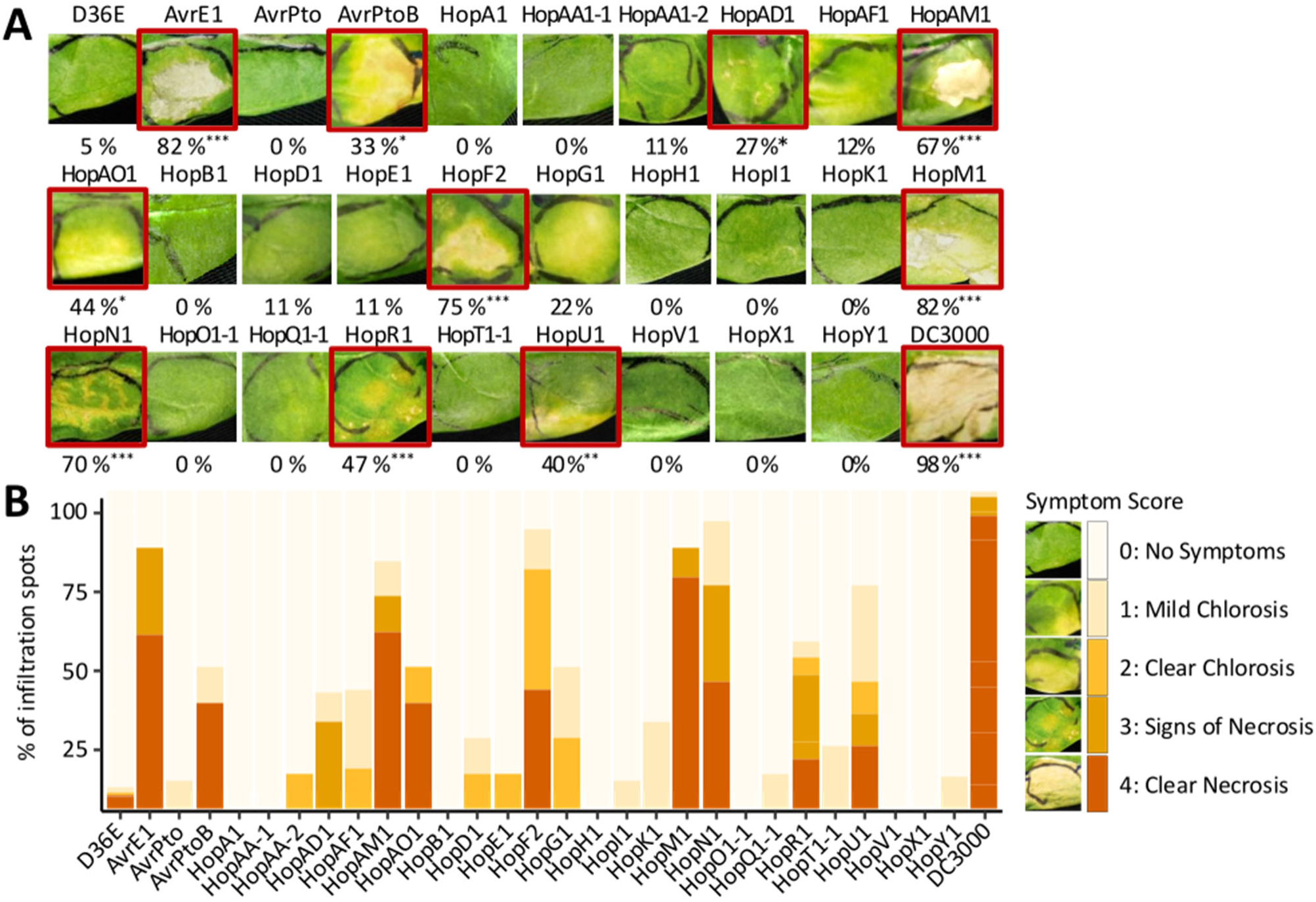
DC3000 effectors in D36E can reconstitute DC3000-like symptoms in spinach. **A)** Representative images of spinach leaves 6–7 days post-infiltration (dpi) with D36E, DC3000, and D36E strains expressing individual *P. syringae* DC3000 effectors. Symptoms were scored using a symptom scoring matrix ranging from 0 (no symptoms) to 4 (clear necrosis), with additional examples in Figure S2. The percentages below each image indicate the proportion of infiltration sites that developed visible symptoms (symptom score >1). D36E, serving as a negative control, induced symptoms at a negligible rate (5%), while DC3000 caused severe symptoms in most infiltration sites (98%). Effectors that significantly influenced symptom development (*p* < 0.05) are highlighted with a red border. Statistical significance was assessed using a Fisher’s Exact test (*** *p* < 0.001, ** *p* < 0.01, * *p* < 0.05). **B)** Stacked bar graph showing the frequency distribution of symptom scores for each tested effector, using a colour gradient from light orange (mild chlorosis, symptom score 1) to dark orange (clear necrosis, symptom score 4). D36E bacteria expressing individual effectors, along with controls (D36E and DC3000), were infiltrated at OD_600_ = 0.4, with symptom development assessed at 6–7 dpi (*n* = 8–12 infiltration sites per treatment).

### *AvrE1*, *HopM1* and *HopU1* significantly affect bacterial proliferation in spinach

To assess whether D36E effector delivery can identify effectors that enhance bacterial proliferation, we measured the contribution of 28 single effectors on early D36E growth in spinach (Figure 3A). Bacterial titres at 2 dpi revealed three effectors that significantly affected D36E growth. Among them, AvrE1 and HopM1 significantly increased D36E titres by a fold difference of log_10_ 2.3 and 1.3 respectively, whereas HopU1 reduced them by a fold difference of log_10_ 0.9 (Figure 3B). The remaining D36E strains showed no significant differences in titres compared to the D36E control, suggesting a limited impact on early proliferation. Notably, while *HopN1*, *HopAM1*, and *HopF2* induced symptoms similar to those of DC3000, they did not enhance D36E proliferation (Figures 2, 3B). Only *AvrE1* and *HopM1*, showed a correlation between effector-driven necrosis and bacterial proliferation, suggesting that they actively manipulate the host to create a beneficial niche.

**Figure 3:**
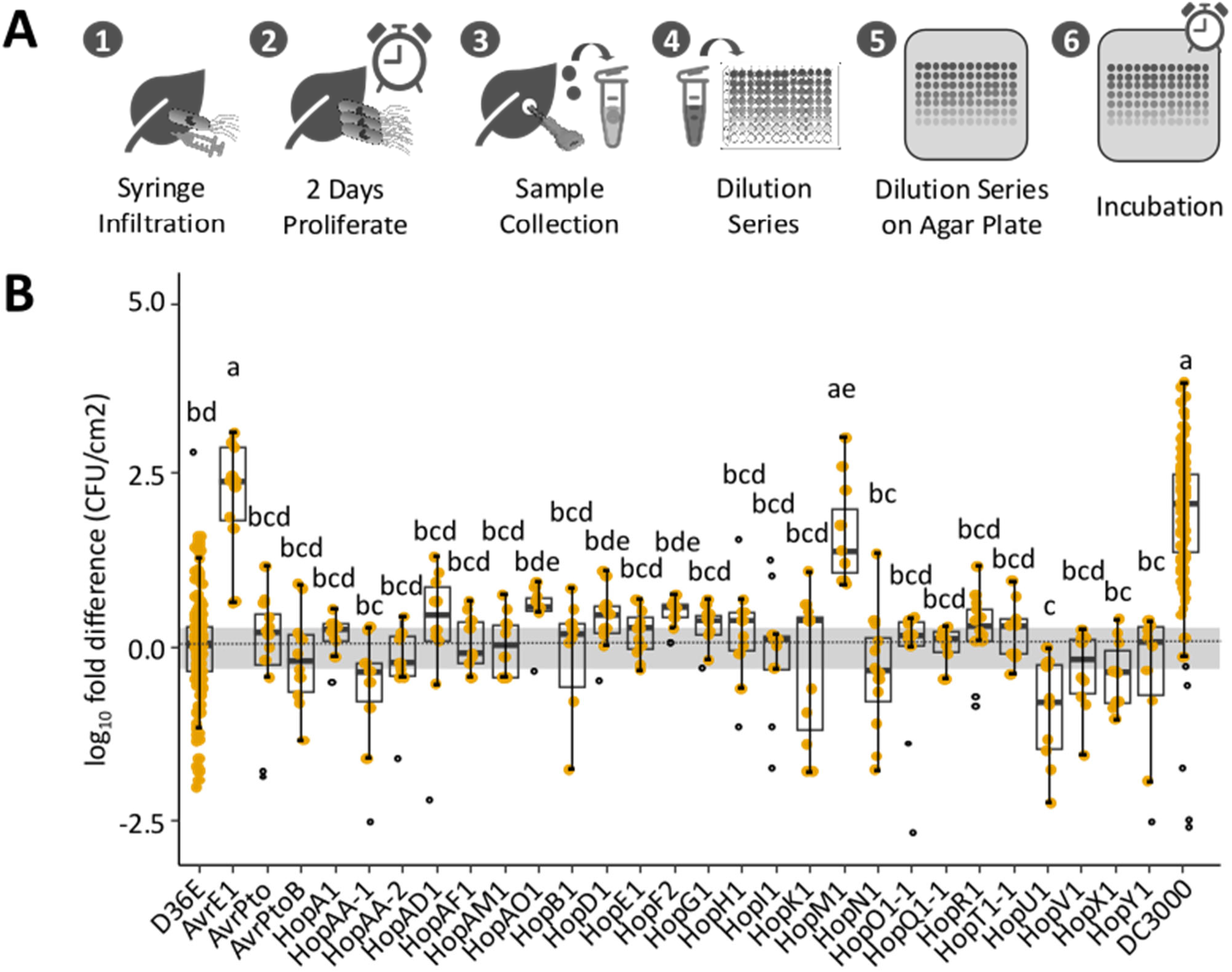
Expression of *AvrE1*, *HopM2*, and *HopU1* significantly alters D36E proliferation in spinach. **A)** Schematic overview of the bacterial quantification workflow. Spinach leaves were syringe-infiltrated with bacterial suspensions (Step 1), followed by a 2-day proliferation period (Step 2). Samples were then collected (Step 3) and subjected to serial dilution (Step 4) before plating on agar (Step 5) and incubating for colony counting (Step 6). **B)** Effect of individual DC3000 effectors on D36E proliferation, measured as the log_10_ fold difference in colony-forming units per square centimetre (CFU/cm^2^) relative to the D36E negative control at 2 days post-infiltration (dpi). Each point represents an individual infiltration site, with measurements based on three pooled leaf discs (*n* = 8–12 per treatment). Statistical significance was determined using one-way ANOVA followed by Tukey’s Honestly Significant Difference (HSD) test. Different letters above boxplots indicate statistically significant differences (*p* < 0.05).

### Inoculum density influences effector-driven phenotypes in spinach

To further investigate the relationship between bacterial proliferation and necrotic symptoms, we conducted infiltration experiments using 10-fold and 100-fold lower inoculum densities (OD_600_ equal to 0.04 and 0.004) for a subset of bacterial strains (Figures S3A-B). We focused on the control strains D36E and DC3000, as well as D36E carrying two effectors with contrasting effects: *AvrE1*, which enhanced both symptom development and bacterial proliferation, and *AvrPto*, which behaved similarly to the D36E control. Additionally, we measured bacterial titres at 6 dpi, when visual symptoms were previously assessed (Figure S3C).

Based on prior assays, we expected D36E and *AvrPto* populations to decline between 0 dpi and 2 dpi, while DC3000 and *AvrE1* titres were anticipated to increase (Figures 2, 3), and as bacterial titres remained stable up to 5 dpi (Figure S1), we expected similar trends at 6 dpi. Overall, bacterial titres scaled proportionally with reduced inoculum densities, except for *AvrE1* at 100-fold dilution, which no longer exhibited enhanced growth (Figure S3A). Interestingly, when diluted 10-fold, DC3000 exhibited a greater increase in growth (two orders of magnitude) compared to the original OD_600_ of 0.4, which showed less than a one-order-of-magnitude increase, suggesting that bacterial growth may plateau at higher inoculum densities. Regardless of inoculum density, DC3000 consistently triggered strong necrosis, whereas *AvrE1*-induced necrosis was markedly reduced at lower densities (Figure S3B). These results indicate that assay outcomes can be dose-dependent, with a minimum inoculum density potentially required to detect relevant phenotypes in spinach using the D36E effector delivery system. At 6 dpi (OD_600_ equal to 0.4), titres of D36E and *AvrPto* were lower than at 2 dpi, whereas *AvrE1* titres remained stable (Figure S3C). Notably, DC3000 titres significantly decreased between 2 dpi and 6 dpi (Figure S3C), suggesting that bacteria in necrotic infiltration sites ultimately die.

### *AvrE1*, *HopM1* and *HopAD1* reduce MAMP-induced ROS bursts in spinach

Reactive oxygen species (ROS) bursts are an early immune response in plants, acting as a defence against plant pathogens (Ngou et al., 2022). Consequently, successful pathogens have evolved mechanisms to suppress ROS bursts (Jwa and Hwang, 2017). To investigate the role of bacterial effectors in modulating early immune responses in spinach, we adapted a ROS burst assay for D36E-based effector screening, following established *Nicotiana benthamiana* methodologies (Johanndrees et al., 2022; Figure 4A).

**Figure 4:**
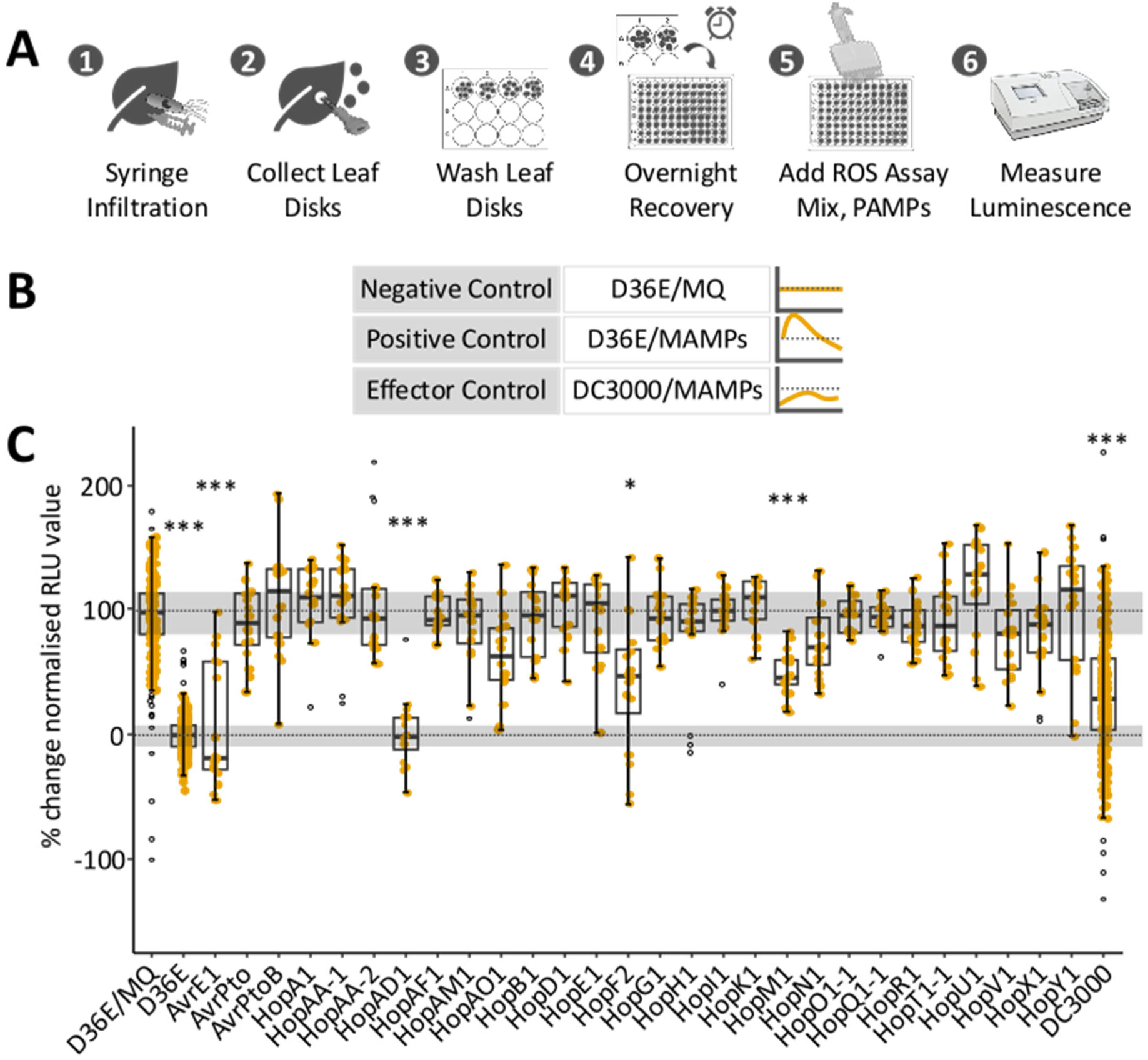
*AvrE1*, *HopAD1*, and *HopM1* can suppress the ROS burst in spinach when delivered by D36E. **A)** Overview of the experimental procedure used to analyse reactive oxygen species (ROS) bursts in spinach. Leaves were pre-infiltrated with D36E strains carrying individual effectors, followed by ROS burst elicitation using a MAMP mix. Chemiluminescence in relative light units (RLUs) was recorded as a measure of ROS production. **B)** Description of controls used in ROS burst assays: a negative control (D36E/MQ) in which leaves are pre-infiltrated with effectorless D36E and mock-elicited, a positive control (D36E/MAMPs) in which a ROS burst is triggered with a MAMP-mix derived from heat-lysed D36E bacterial cells in leaves pre-infiltrated with effectorless D36E, and a suppressed control (DC3000/MAMPs) in which a ROS burst is suppressed in leaves pre-infiltrated with DC3000. **C)** Quantification of ROS burst suppression following infiltration of D36E expressing 28 individual DC3000 effectors. The percent change in ROS burst was calculated relative to the negative control. One-way ANOVA followed by Dunnett’s post-hoc test was applied to compare normalized and scaled RLU of D36E effector strains to the negative control. Asterisks indicate significant differences (*p* < 0.001, *p* < 0.01, *p* < 0.05; *n* = 16 leaf discs per treatment).

First, we screened known ROS burst elicitors to identify the most potent triggers in spinach leaf discs. We tested chitin (Sánchez-Vallet et al., 2015), the flg22 peptide (Mersmann et al., 2010), the nlp24 peptide from a downy mildew Nep1-like protein (Oome et al., 2014), heat-lysed *P. effusa race 1* sporangia (Pe1 sporangia) (Klein et al., 2020; Skiadas et al., 2022), and a microbe-associated molecular pattern (MAMP) extract from heat-lysed D36E cells (Wei et al., 2015) (Table S5). Among these, chitin, the MAMP-extract, and Pe1 sporangia triggered significant ROS bursts, with the MAMP-extract inducing the strongest response (Figure S4). Therefore, we selected the MAMP-extract as elicitor for subsequent assays.

Next, we established controls for D36E-compatible ROS assay. We compared ROS production in spinach leaf discs pre-infiltrated with either D36E or a mock solution (Figure S5). Pre-infiltration with D36E alone (D36E/MQ) did not alter ROS levels compared to the mock control, whereas treatment with the MAMP-extract after D36E infiltration (D36E/MAMPs) triggered a robust ROS burst. Based on these results, we used D36E/MQ as the negative control and D36E/MAMPs as the positive control (Figure 4B). When we tested DC3000 (OD_600_ equal to 0.4), we observed significant ROS suppression compared to the D36E/MQ negative control, which we attributed to effector-mediated suppression (Figure S6).

Finally, we screened the D36E single-effector library to identify effectors that suppress the MAMP-extract-induced ROS burst in spinach (Figure 4C). Among the 28 tested effectors, *AvrE1*, *HopAD1*, and *HopM1* significantly reduced ROS production. Notably, *HopAD1* suppressed ROS bursts to levels comparable to the D36E/MQ negative control, even exceeding the suppression observed in DC3000-infiltrated leaf discs. Most other effectors did not significantly alter ROS burst levels, indicating that the ability to suppress early immune responses is limited to a specific subset of effectors.

### Only AvrE1 and HopM1 largely restored DC3000-like characteristics on spinach

To identify similarities among individual DC3000 effectors across different assays in spinach, we systematically compared phenotypic outcomes for each strain. To enable a direct comparison, we standardized phenotypic responses using Z-scaling and applied hierarchical clustering (Figure 5). This analysis revealed five distinct effector clusters (Clusters I–V), each characterized by a unique combination of phenotypic patterns. Cluster I includes DC3000, *HopM1*, and *AvrE1*, confirming that these two effectors can largely restore DC3000-like effects for the assessed phenotypes in spinach. Cluster II consists of a single strain, *HopAD1*, which strongly suppressed the ROS burst while not affecting proliferation and causing symptoms in 27% of infiltration spots (Figures 2-5), suggesting that ROS burst suppression alone is insufficient to restore D36E proliferation. Cluster III includes *HopR1*, *HopAM1*, *HopN1*, *HopF2* and *HopAO1* which induced significant symptoms, while not altering ROS bursts or bacterial proliferation (Figures 2-5). Cluster IV comprised *AvrPtoB* and *HopU1*. These effectors reduced bacterial proliferation (significantly for *HopU1*, Figure 3B), but had no impact on ROS production or symptom development (Figures 2, 4C). Cluster V includes the remaining 18 effectors thatdid not show any detectable phenotypic responses in our assays.

**Figure 5:**
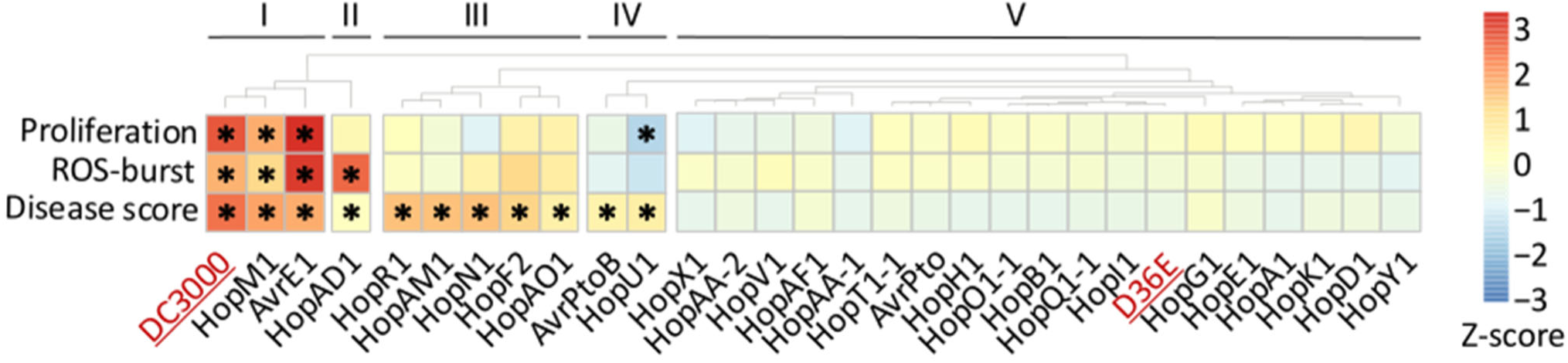
Systematic analysis of the functional effector screens highlights that D36E expressing *AvrE1* and *HopM1* restore DC3000-like phenotypes on spinach. Assay read-outs were Z-scaled to enable direct comparison of phenotypic outcomes across 28 individual DC3000 effectors expressed in D36E. Hierarchical clustering of Z-scaled values identified five distinct clusters (Cluster I – Cluster V), reflecting different phenotypic patterns. The largest group of 18 DC3000 effectors (Cluster V), did not significantly affect spinach in any of the assays, while *AvrE1* and *HopM1* appear as the most important contributors to DC3000-like symptoms (Cluster I). Asterisks (*) indicate significant effects (*p* < 0.05).

### Consistent responses across cultivars supports broad applicability of type 3 effector delivery in spinach

Consistent phenotypic responses of the D36E-based delivery assay spinach cultivars are important for future applications. We therefore assessed the robustness of our assay setup by testing multiple spinach cultivars for their responses to key controls (D36E and DC3000) (Figure S7). We infiltrated five spinach cultivars, i.e., Whale, Hydrus, NL3, NIL5 and NIL6 (OD_600_ equal to 0.4), and found that these all responded similarly to the responses previously recorded in Viroflay, exhibiting necrosis with DC3000 infiltration and no symptoms with D36E (Figure S7A). Similarly, bacterial proliferation patterns were consistent across cultivars. Like Viroflay, D36E titres decreased between 0 dpi and 2 dpi, whereas DC3000 titres increased in cultivars Whale, NIL3 and Caladonia (Figure S7B). To assess the suitability of our ROS assay setup, we tested D36E/MQ (negative control), D36E/MAMPs (positive control) and DC3000/MAMPs (suppressed control) on Whale and NIL3. The responses of these cultivars were comparable to those observed in Viroflay (Figure S7C). Together, these findings indicate that our newly established spinach type 3 delivery assays can be reliably applied across different spinach cultivars.

## Discussion

To advance knowledge-driven resistance breeding in spinach functional studies on pathogen effectors are of high importance. Here, we established an *in-planta* effector screening system using the *Pseudomonas syringae* pv. *tomato* DC3000 polymutant D36E (D36E) as a protein delivery system. This system allowed us to systematically assess the contribution of DC3000 effectors to disease susceptibility and symptom development in spinach. Among the 28 bacterial effectors tested, we identified *AvrE1* and *HopM1* as key drivers of the DC3000 phenotype, both promoting necrosis and bacterial proliferation, while also suppressing the reactive oxygen species (ROS) burst. We also identified *HopAD1* as a potent suppressor of ROS bursts, though its effect on bacterial fitness in spinach remained inconclusive. Moreover, we found that expression of the effectors *HopAM1*, *HopAO1*, *HopF2*, *HopN1* and *HopR1* contributed to symptom development without significantly affecting D36E proliferation. To our knowledge, this represents the first report of a functional *in planta* effector screen in spinach, establishing a foundation for future resistance studies.

### AvrE1 and HopM1 drive bacterial virulence in spinach

A major challenge in interpreting our findings is the lack of a confirmed, recognized effector in spinach to benchmark our observations. In the non-host plant *Nicotiana benthamiana,* AvrE1, HopM1 and HopAD1 act as elicitors of effector-triggered immunity (ETI), inducing a characteristic response of programmed cell death and restricted bacterial growth (Wei, Zhang et al. 2018). In spinach, we found that *AvrE1* and *HopM1* induce necrosis similar to DC3000. Unlike observations in *N. benthamiana*, necrosis in spinach was accompanied by increased bacterial titers compared to the negative control D36E (Figures 2, 3). We initially attributed this to relatively high infiltration density. However, the increase in bacterial titres between 0 dpi and 2 dpi persisted even at lower inoculum densities and at later timepoints (Figures S3A, C). Therefore, we hypothesise that necrosis is the result of successful bacterial infection rather than an active plant immune response.

Expression of *AvrE1* and *HopM1* in D36E resulted in comparable phenotypic outcomes in spinach (Figure 5). Functional redundancy among these two effectors was extensively described in model plants including DC3000’s natural host, tomato, but also in *N. benthamiana* and *Arabidopsis thaliana* (*Arabidopsis*) (DebRoy et al., 2004; Badel et al., 2006; Cunnac et al., 2011). In DC3000, these effectors are encoded alongside each other at the conserved effector locus (*CEL*) that is widely distributed among plant-infecting bacteria of the genus *Pseudomonas* (Baltrus et al., 2011). Both effectors contribute to the creation of an aqueous micro-environment on host plants, which is crucial for successful *Pseudomonas* infection (Roussin-Léveillée et al., 2022). In spinach, we observe water-soaked lesions—symptomatic for the establishment of an aqueous micro-environment—following inoculation with D36E bacteria carrying *AvrE1* or *HopM1,* suggesting that their virulence targets are likely conserved in spinach.

On the molecular level, AvrE1 and HopM1 are distinct, belonging to different molecular families. AvrE1 belongs to a broadly conserved AvrE1/DspA/E/HopR1 superfamily of effectors (Kvitko et al., 2009; Baltrus et al., 2011), while HopM1 is part of a less conserved effector family (Baltrus et al., 2011; Jayaraman et al., 2020). In its superfamily, AvrE1 is the best characterized effector, recently demonstrated to induce water-soaked lesions by forming a pore-forming beta barrel, thereby creating a permeable membrane channel (Nomura et al., 2023). The pore-forming function appears not to be shared by HopM1, which does not exhibit the distinct beta barrel structure or other prominent molecular features (Nomura et al., 2023). In spinach, we observed that *AvrE1*, more than *HopM1*, could induce DC3000-like symptoms by D36E (Figure 5A), possibly via its pore-forming function.

HopM1, like AvrE1, has been implicated in creating water-soaked lesions to enhance virulence in *Arabidopsis* (Xin et al., 2016), which aligns with early phenotypes we observed in spinach. HopM1 was shown to target the proteasome, a central point of regulation in plant immunity (Üstün et al., 2016). HopM1 enhances AtMIN7 proteasome-dependent degradation consequently interfering with vesicle transport in *Arabidopsis* (Nomura et al., 2006; Xin et al., 2016). In addition, HopM1 destabilises a range of other proteins in a proteasome-dependent manner that include, e.g., 14-3-3 proteins like GRF8/TFT1 (Lozano-Durán et al., 2014). HopM1 was shown to reduce PAMP-triggered ROS bursts, similar to chemical destabilisation of 14-3-3 proteins or a targeted silencing of GRF8/TFT1, suggesting that HopM1 targets 14-3-3 proteins to supress early immune responses like ROS bursts (Lozano-Durán et al., 2014). We observed that infiltration of spinach leaves with D36E carrying *HopM1* significantly suppressed the ROS burst, suggesting that the interference with 14-3-3 proteins might be conserved (Figure 4C).

### HopAD1 suppresses the ROS burst but does not enhance bacterial growth

Infiltration of spinach leaves with D36E bacteria carrying *HopAD1* led to a strong suppression of the ROS burst and the development of mild necrosis in 27% of infiltration spots (Figures 2, 4C). Compared to AvrE1/HopM1, less information is available on HopAD1, as it was initially dismissed as a non-functional effector (Chang et al., 2005). However, Wei and colleagues (2015) already described its potential to suppress the ROS burst and trigger cell death in *N. benthamiana*. When the effector AvrPtoB is present, like in the case of DC3000, HopAD1’s immunogenic activities are masked in *N. benthamiana* leading to the earlier dismissal of this effector (Wei et al., 2015). Since the nearly effector-less DC3000 mutant D28E does not encode *AvrPtoB*, Wei and colleagues could successfully reveal its function (Wei et al., 2015).

While in spinach and in *N. benthamiana*, infiltration by D36E bacteria carrying *HopAD1* yielded strong suppression of the ROS burst, only in *N. benthamiana*, infiltration at high titres also resulted in consistent necrotic symptoms (Wei, Zhang et al. 2018, Figures 2, 4C). In *N. benthamiana*, cell death did not necessarily lead to altered virulence, mimicking our findings in spinach (Wei, Zhang et al. 2018, Figures 2, 5). We propose that the suppression of the MAMP-induced ROS burst is a primary function of HopAD1, which by itself appears to be insufficient to promote bacterial proliferation. Despite the identification of HopAD1’s immunogenic activity, the molecular mechanisms underlying its ability to suppress the ROS burst while leaving bacterial proliferation unaffected in spinach remain elusive.

### Effector redundancy and synergy in spinach

In nature, effectors often act synergistically and display redundancy. For instance, our findings reveal that the Δ*CEL* mutant, characterized by the absence of *AvrE1*, *HopM1*, *HopN1* and *HopAA-1* (Alfano et al., 2000; Wei and Collmer, 2018), induced robust visible symptoms and exhibited proliferation similar to DC3000 at 2 dpi on spinach (Figure 1). This finding contrasts with our observations of single effectors, wherein only *AvrE1* and *HopM1* could significantly enhance proliferation of D36E bacteria on spinach. However, its DC3000-like virulence underscores the importance of synergistic effector action in nature. The combined influence of the remaining effectors in the Δ*CEL* mutant appears to compensate for the absence of *AvrE1* and *HopM1* in spinach, emphasizing the intricate synergies among effectors in shaping the overall pathogenic response. Evidence for genetic interactions between DC3000 effectors has been previously found in *N. benthamiana*, where HopAD1-dependent disease symptoms are suppressed by AvrPtoB, and HopI1 suppressed cell death elicited by AvrE1, HopM1, HopR1, HopQ1-1 and HopAM1 (Wei et al., 2015).

## Conclusion

We present the first report of a functional *in planta* effector screen in spinach. In the future, the D36E delivery system can be harnessed for characterising spinach pathogen effectors to identify those recognised by the spinach immune system. Compared to native spinach pathogens such as the obligate biotrophic oomycete *P. effusa*, the D36E effector delivery provides a simplified system to study effectors in their native hosts. In contrast to *Agrobacterium*-mediated transient expression, our data suggest that the D36E system performs similarly across spinach cultivars and does not require extensive fine-tuning for biochemically distinct effectors like AvrE1, HopM1 or HopAD1 (Figures 2, S7) (Zhang et al., 2024). Thus, we here introduce a novel approach for functional effector studies in spinach with potential for high-throughput screening.

## Material and Methods

### Plant material

All *in planta* assays were conducted using the spinach cultivar Viroflay (Pop Vriend Seeds, The Netherlands). Seeds were sown and plants were cultivated on Primasta® potting soil within a plant growth room providing a stable environment (21 °C, 16 hr photoperiod, 70% relative humidity, 200 µmol m ^2^ s^-1^ light intensity). Assays were initiated when the first true leaves were fully expanded, and the second set of true leaves started development, typically occurring between 2.5 and 3 weeks post sowing. To ensure consistency, phenotypically-similar plants were selected for all experiments.

### *Pseudomonas* infiltration

To identify a suitable bacterium for T3SS-dependent effector delivery into spinach, we tested different *Pseudomonas* strains. Preparation of bacterial cultures and infiltration was performed as previously described with the following adaptations (Wei et al., 2013; Liu et al., 2015). Bacteria were grown for two days on solid King’s B medium (KB) supplemented with antibiotics at 28 °C (King et al., 1954). Antibiotic resistances and their concentrations are listed in Table S2. Bacteria were re-streaked to new KB agar plates 2 days prior to infiltration (28 °C). For infiltration, bacteria were scraped off the KB agar plates and suspended in infiltration medium (10 mM MgSO_4_, 50 µl/l Coatosil 1220 surfactant in MilliQ (MQ) water). Bacteria were infiltrated from the abaxial side of fully expanded spinach leaves with a needleless syringe. Infiltrated spinach plants were placed in trays and covered from the light before transfer to the climate chamber. The cover was removed after one day, exposing the plants to the light conditions. Screening for symptom development and bacterial proliferation were performed as described below.

### Generating the D36E single effector library

For single effector screens in this study, we obtained the DC3000 single effector library in pENTR/SD/D-TOPO backbones and the matching Gateway destination vector pCPP5372 via the non-profit plasmid repository <u>Addgene</u> (Wei et al., 2018). Single *Escherichia coli* colonies were picked and grown in overnight liquid Luria-Bertani (LB) medium supplemented with selective antibiotics (Table S1, S2, S3). Plasmids were isolated using the QIAprep® Spin Miniprep Kit (Qiagen), followed by Sanger sequencing with the M13 forward primers to confirm effector identity (Table S4). We mobilised DC3000 effector genes into *pCPP5372* by Gateway® LR cloning as previously described (Wei et al., 2018). For propagation, plasmids were transformed into Top10 *E. coli* cells and grown on LB agar plates with selective antibiotics (Table S1, S2, S3). We selected positive colonies by colony-check PCR using effector-specific reverse primers and the pCPP5372_pre_attR1 forward primer (Table S4). Effector expression plasmids were mobilised from *E. coli* into D36E by triparental mating using the pRK2013 *E. coli* HB101 with the *pRK2013* helper plasmid (Table S1).

### Visual symptom scoring

To assess the visual development of symptoms, *Pseudomonas* strains were prepared and infiltrated as described above. After infiltration, plants were grown until visual assessment of symptoms at 6-7 dpi. To evaluate the symptoms of the single effector library on spinach, we infiltrated each leaf with the negative control D36E and the positive control DC3000, alongside D36E carrying individual effectors. D36E does not trigger symptoms while DC3000 does. The two remaining infiltrated sites per leaf were infiltrated with the individual effector lines. We established a symptom scoring matrix ranging from symptom score 0 to symptom score 4, corresponding to no symptoms (0), mild chlorosis (1), clear chlorosis (2), signs of necrosis (3) and clear necrosis (4) (Figure 2, S3). A Fisher’s Exact test was applied to compare incidences of symptom-positive infiltration sites (symptom score >1) against the incidence of symptom-free infiltrated sites (symptom score <2). The number of infiltration sites showing a specific score was plotted using the *R* package *ggplot2* (Wickham, 2016; Team;, 2021).

### D36E proliferation assays

To better understand the connection between visual symptom and bacterial titres of *Pseudomonas* strains, we assessed bacterial proliferation as a proxy for virulence by evaluating bacterial titres. We adapted the published protocol by Liu et al. (2015) (Liu et al., 2015). First, spinach leaves were infiltrated with *Pseudomonas* strains as described above. Then, infiltrated leaves were allowed to dry for 1.5 to 2 hours before collecting 0 dpi samples. These samples were used to verify successful, homogenous infiltration. Three leaf discs (4 mm diameter) per infiltration sites were collected in 500 µl infiltration medium containing three glass beads (2 mm). Bacterial cells were extracted from spinach tissue by applying tissue lysis using the TissueLyser II (Qiagen, 85300) until leaf discs were completely homogenized (twice 2 min, 30 hz s^-1^). We prepared 10-fold serial dilutions of homogenized samples in infiltration medium using 96-well plates. From all dilution steps, 5-µl droplets were transferred to square KB agar plates with appropriate antibiotics (Table S1, S2). Droplets were allowed to dry, and plates were then sealed with Parafilm (Bemis, PM996) and incubated for 1.5 to 2 days at 28 °C until colonies were clearly visible. Bacterial titres were estimated by counting colony forming units (CFUs) on KB agar plates. CFUs counts at the highest dilution level with >3 three colonies, were used to calculate the number of CFUs per cm^2^ spinach leaf.

Data were processed in *R* and plotted in *ggplot2* (Wickham, 2016). For statistical analysis of initial *Pseudomonas* screens (Figure 1), we assessed differences in proliferation between 0 dpi and 2 dpi using a two-tailed t-test. For statistical analysis of the single effector library screens at 2 dpi, relative to the D36E negative control, we applied an ANOVA followed by a Tukey post-hoc test. Scripts and raw data are available on <u>GitHub</u>.

### D36E-compatible ROS burst assays in spinach

We modified the ROS burst assay from (Johanndrees et al., 2022), based on (Bisceglia et al., 2015) to make it compatible with the D36E effector delivery system in spinach. First, we identified elicitors for ROS bursts in spinach. Spinach leaf discs (4 mm diameter) were harvested from 2.5-3-week-old plants and washed trice in MQ water (30 min, gentle shaking). Discs were arranged on a 96-well plate (Greiner Bio-One; 655075) in 200 μl MQ water, wrapped in aluminium foil to shield from light, and incubated at 21 °C overnight. MQ was replaced with the luminol analogue L-012 sodium salt (Merck SML2236, final concentration 180 μM) and horseradish peroxidase (Merck; P8125-5KU, 0.125 units per reaction) solution. To each sample well, 50 µL elicitor solution was added at concentrations specified in Table S5. We tested chitin, flg22, nlp24 and heat-lysed *P. effusa* race-1 sporangia (15 min, 100 °C; *Pe1* sporangia) and heat-lysed D36E cells (15 min, 100 °C; MAMP-mix). Additional details on elicitors can be found in Table S5. Chemiluminescence was measured every 2.5 min in a GloMax® 96 Microplate Luminometer (Promega).

To investigate the impact of effectors on the ROS burst in spinach using the D36E effector delivery system, we first established the proper controls. Bacteria were cultivated and infiltrated at 0 dpi as described above. Infiltrated spinach plants were dried for 1.5 to 2 hours before proceeding with the ROS assay protocol as described above. D36E/MQ was established as negative control, D36E/MAMPs as positive control, DC3000/MAMPs as effector-positive control which suppresses ROS bursts (Figure S5, S6, S7). To compare ROS burst assays, log_2_ of raw Relative Light Units (RLUs) were calculated and plotted using *ggplot2* in R (Wickham, 2016). Additional features, time to peak ROS scores, peak ROS score and total ROS scores, were calculated using Python 3.10 (van Rossum and Drake, 2009). To screen the complete single effector library for alterations in MAMP-mix elicited ROS bursts, all three controls were included. To analyse the variable results of single effector library screens, we processed, normalised and transformed raw RLUs in Python 3.10 with *pandas* version 1.5 using the internal positive and negative controls for scaling (van Rossum and Drake, 2009; team, 2020). Statistical analysis using ANOVA followed by a Dunnett’s post hoc test and visualization was conducted in *R* using *ggplot2* (Wickham, 2016; Team;, 2021). All scripts and raw data available on <u>GitHub</u>.

### Systematic analysis of single effector library screens on spinach

We constructed a heatmap using the *pheatmap* package in *R* (Kolde, 2018; Team;, 2021). Phenotypic outcomes are presented by extraction of representative single-values from visual symptom scoring, bacterial proliferation assays and ROS burst assays. ROS burst values were inverted by multiplication with -1 to allow for intuitive presentation of effector contribution to symptoms. We clustered rows using the Euclidean distance metric combined with complete linkage mapping for hierarchical clustering (*pheatmap* default). To facilitate an ordered discussion of the heatmap, we manually rotated the leaves of the dendrogram. Linear regression was conducted by calculating *R^2^* values using a linear model, corresponding p-values are presented. Results were plotted in *R* using *ggplot2* (Wickham, 2016; Team;, 2021). Scripts and raw data are available on <u>GitHub</u>.

## Acknowledgments

This project was financially supported by the TopSector TKI Tuinbouw & Uitgangsmaterialen, the Netherlands, and four private spinach breeding companies: Enza Zaden Research & Development B.V. (NL), Pop Vriend Seeds (subsidiary of KWS Vegetables BV) (NL), Rijk Zwaan Breeding B.V. (NL), and Syngenta (NL).

We are grateful for the generosity of the research community in sharing bacterial strains. Special thanks to Alan Collmer, Professor Emeritus of Cornell University, and Wei Zhang, Cornell University, for providing the DC3000 D36E and D28E strains, and to Jeff Chang of Oregon State University for sharing the EtHAn strain.

## Supplementary Material

**Figure S1:**
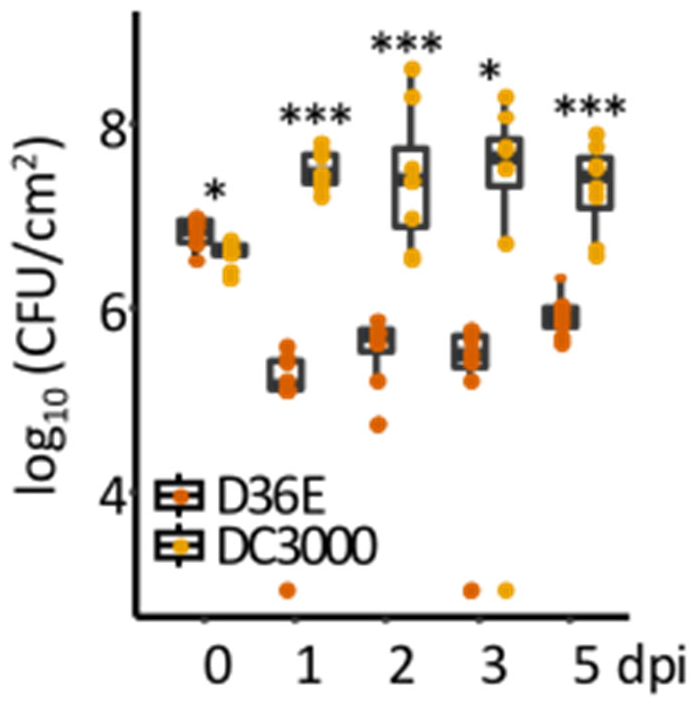
DC3000 and D36E titres stabilize at different levels between 1- and 5-day post-infiltration (dpi) in spinach. To evaluate whether bacterial populations stabilise, we infiltrated bacterial cultures at 0 days post-infiltration (dpi) at OD_600_ equal to 0.4 and tracked titres (colony-forming units or CFU) over five days, sampling 0-3 dpi and 5 dpi. The low variance at 0 dpi is indicative of proper technical reproducibility. Already at 1 dpi it becomes clear that DC3000 cells actively proliferate in spinach while the titres of D36E decrease. Afterwards, both populations appear to stabilize and the difference between titres remains significant until the end of the experiment. For statistical analysis, we applied a two-tailed t-test (*** *p* < 0.001, ** *p* < 0.01, * *p* < 0.05).

**Figure S2:**
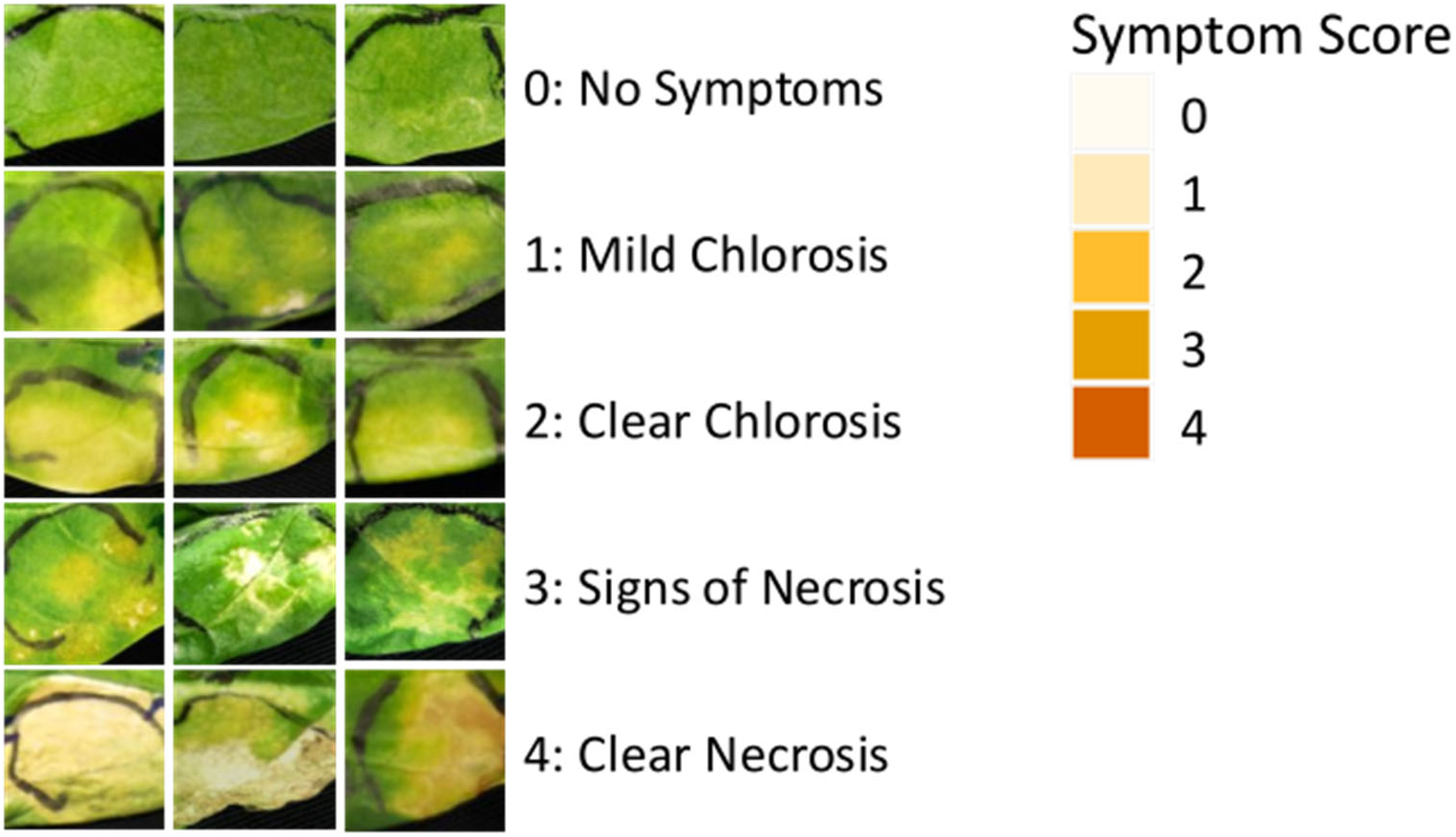
Variation in visual symptoms and symptom scores. We observed substantial variation in symptoms on spinach following infiltration with *Pseudomonas* strains. To categorise visual symptoms and capture the variation of symptoms, we devised a scoring matrix (Figure 2).

**Figure S3:**
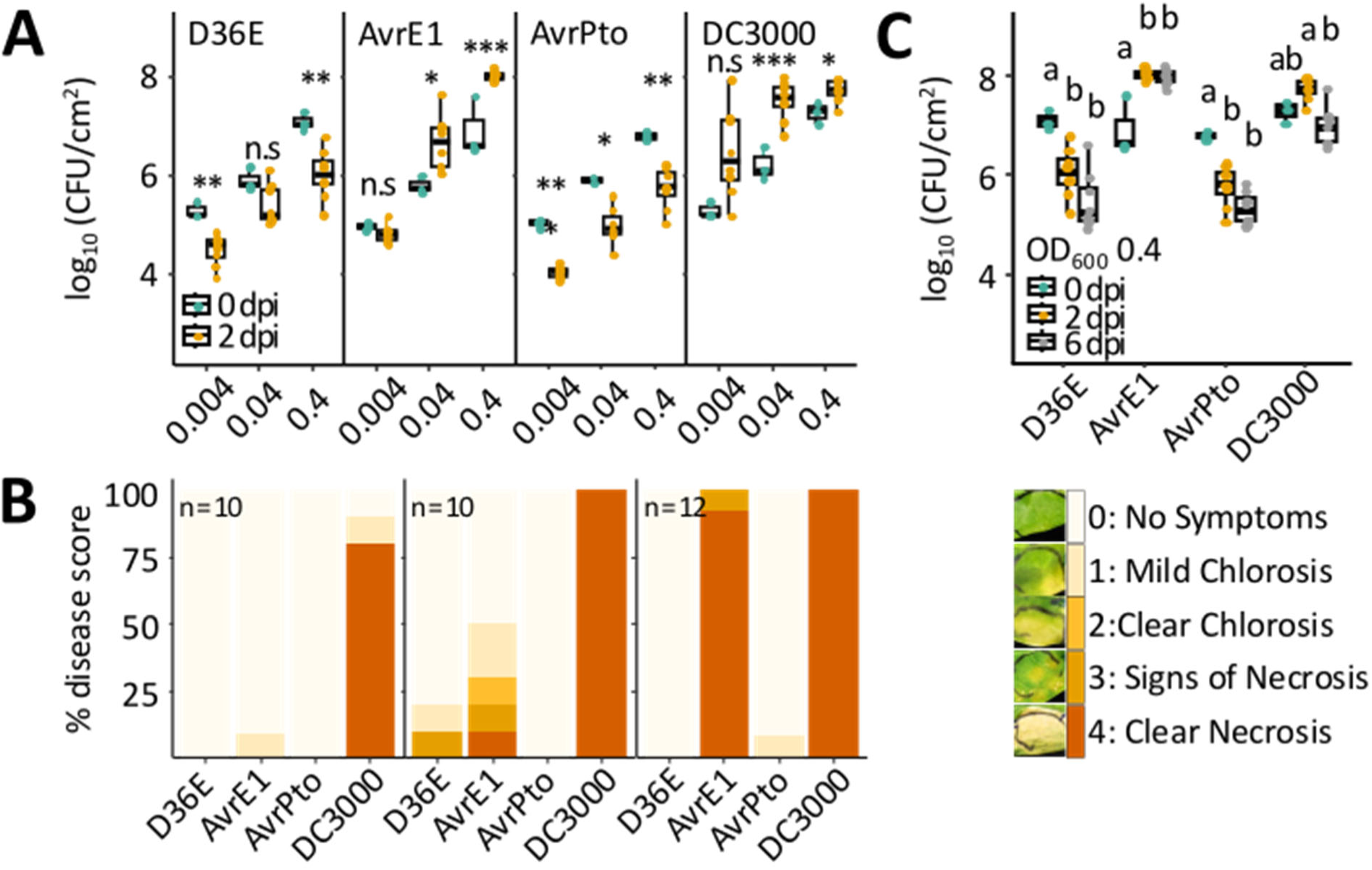
*Pseudomonas* titres and symptom development in spinach at varying inoculum densities and late timepoints. **A)** Bacterial titres develop similarly between 0-day post-infiltration (dpi) and 2 dpi across different inoculum densities. At OD_600_ equal to 0.004, *AvrE1* titres no longer increase between 0 dpi and 2 dpi. In contrast, DC3000 titres increase less than 1 order of magnitude at OD_600_ of 0.4 but increase around 2 orders of magnitude at OD_600_ of 0.04. **B)** Development of symptoms after infiltration of *Pseudomonas* strains at different inoculum densities at 6 dpi (left: OD_600_ equal to 0.004, middle: OD_600_ equal to 0.04, right: OD_600_ equal to 0.4). We scored symptoms in each infiltration site based on the presented symptom scoring matrix ranging from 0 (no symptoms) to 4 (necrotic symptoms). DC3000 causes severe, level 4 symptoms in most infiltration sites at all tested inoculum densities, while *AvrE1* triggered symptoms drop considerably at the two lowest inoculum densities. D36E and *AvrPto* do not cause visible symptoms. **C)** *Pseudomonas* titres at 0 dpi, 2 dpi and 6 dpi following infiltration at OD_600_ equal to 0.4. Titres of D36E and *AvrPto* decrease significantly over time, whereas *AvrE1* and DC3000 titres increase initially, but drop between 2 dpi and 6 dpi in the case of DC3000. We conducted a one-way ANOVA followed by Tukey’s Honestly Significant Difference (HSD) test separately for each strain. Statistical differences are indicated by letters above the boxplots, with n = 3 infiltration sites per treatment.

**Figure S4:**
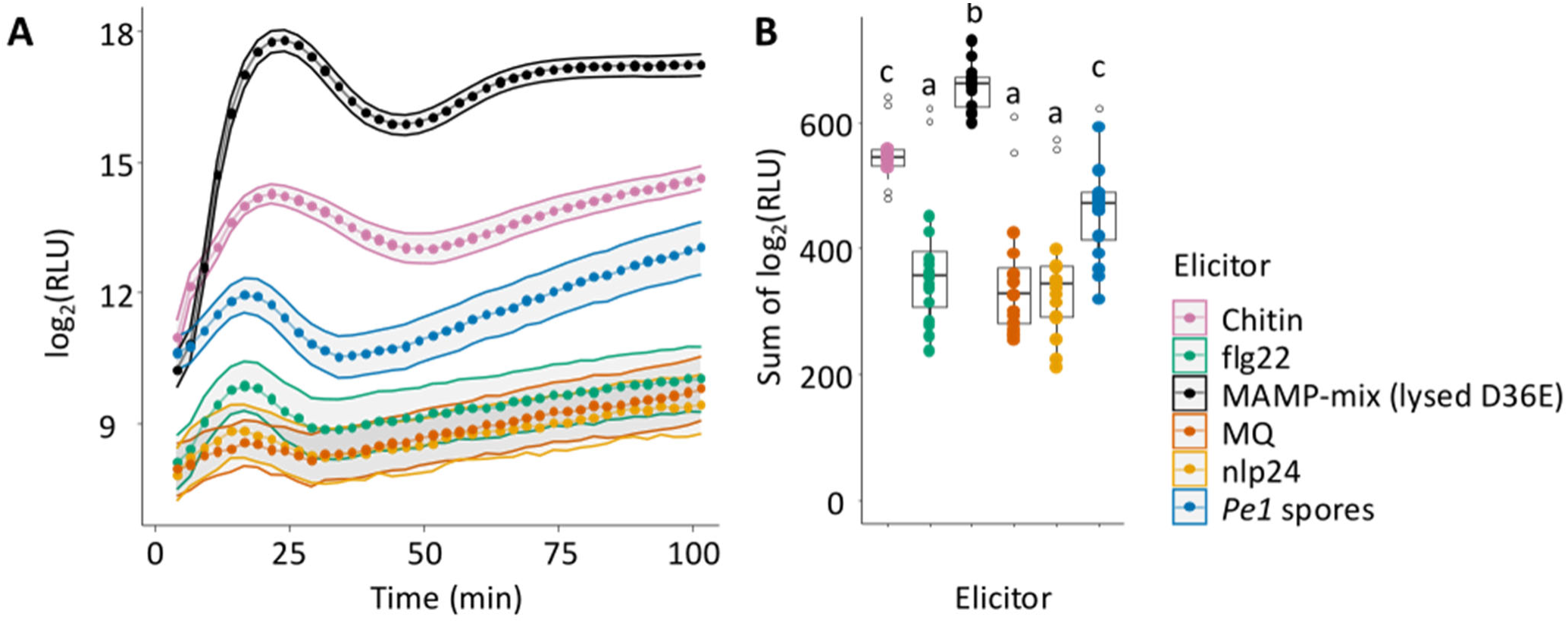
Screening putative reactive oxygen species (ROS) burst elicitors on spinach. We tested five different elicitors for their ability to trigger a rapid ROS burst in spinach, measured as relative light units (**RLU**). We chose the well-described MAMP-type elicitors chitin, flg22 and nlp24, and we included two elicitor mixes, i.e.: heat-lysed D36E (MAMP-mix/MAMP) cells and heat-lysed *P. effusa race 1* (*Pe1*) sporangia. We show results as a log_2_ transformed raw relative light units (RLUs) values (**A**), and the sum of log_2_ transformed raw RLU values over the duration of the experiment (**B**). The MAMP-mix elicited the strongest ROS burst (black), followed by chitin (pink), scoring in sum around 100 log_2_(RLU) lower.

**Figure S5:**
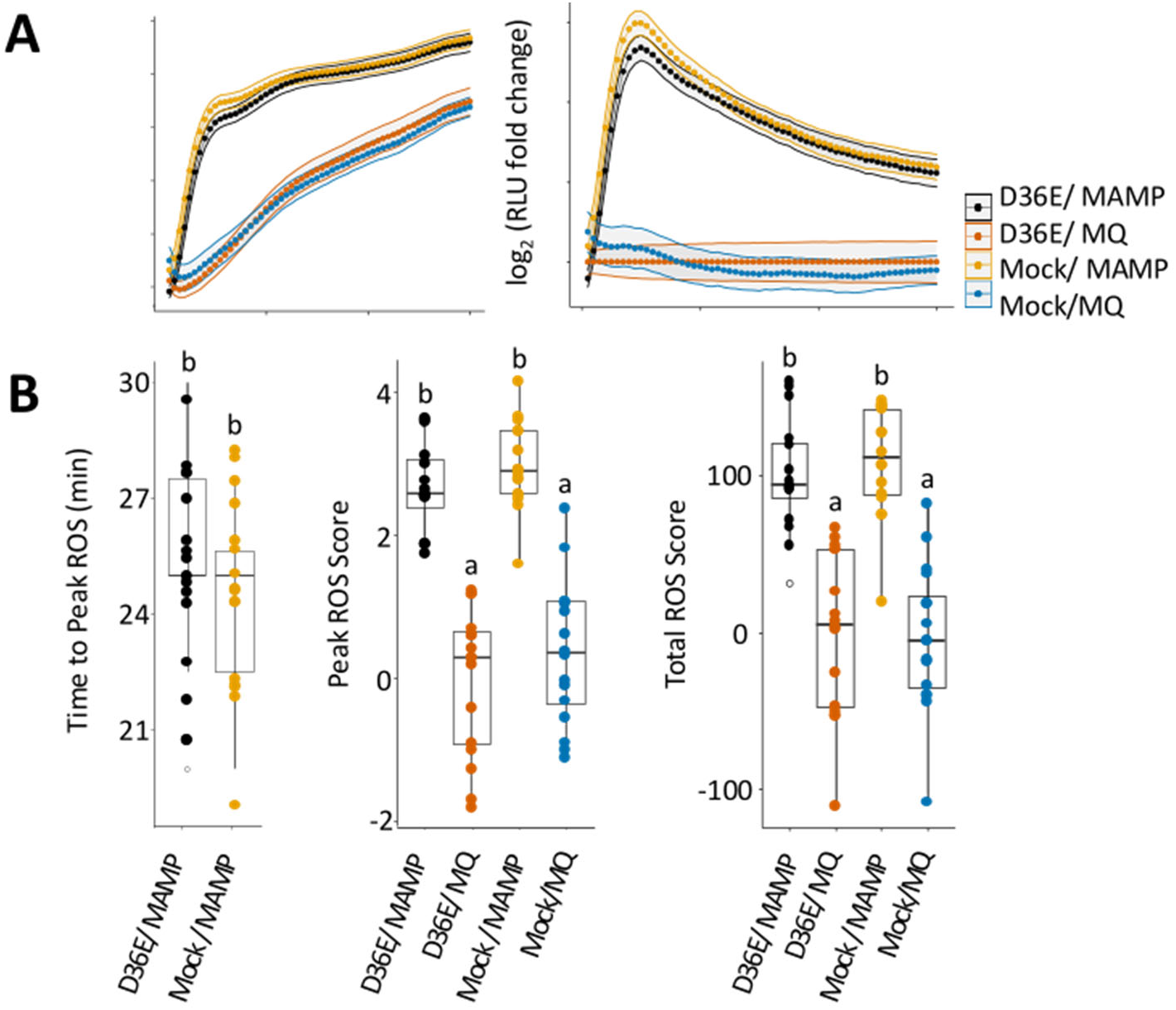
Establishing negative and positive controls for reactive oxygen species (ROS) burst assays on spinach in combination with the effectorless *Pseudomonas syringae pv*. *tomato* strain (D36E) delivery. **A)** To establish ROS burst negative and positive controls, we measured and compared ROS in relative light units (RLU) mock-infiltrated and non-elicited (mock/MQ) and mock-infiltrated and elicited (mock/MAMP) samples with D36E-infiltrated and non-elicited (D36E/MQ) and D36E-infiltrated and elicited (D36E/MAMP) samples. In both comparisons, D36E had no significant impact on ROS burst results, establishing D36E/MQ as a negative control and D36E/MAMPs as a positive control. For all treatments, we noted a consistent rise in ROS production over time, even in the absence of elicitation (**left**). We normalized our data for this steady increase in background ROS levels by calculating the fold change between the negative control D36E/MQ and measurements of other samples at each timepoint (measurement/control measurement) (**right**). Furthermore, the analysis revealed that no major changes in ROS production are to be expected after ∼50 min, which we consequentially used as maximum time during effector screening. **B**) Boxplots display three parameters extracted from the time-resolved graphs depicted in **(A)** after transformation and normalization, i.e.: time to peak ROS (time to maximum ROS production) (**left**), peak ROS scores (maximum of measure ROS scores) (**middle**), and total ROS scores (sum of ROS scores over time) (**right**). Time to peak ROS does not show a statistically significant difference between mock-elicited and MAMP-mix-elicited samples. Peak ROS scores and total ROS scores show significant differences between mock-elicited and elicited samples but no difference between mock-infiltrated and D36E infiltrated samples, confirming the choice of controls for further use. We conducted statistical analysis using ANOVA, followed by Tukey’s Honestly Significant Difference test (*p* < 0.05). Distinct letters denote significantly different groups.

**Figure S6:**
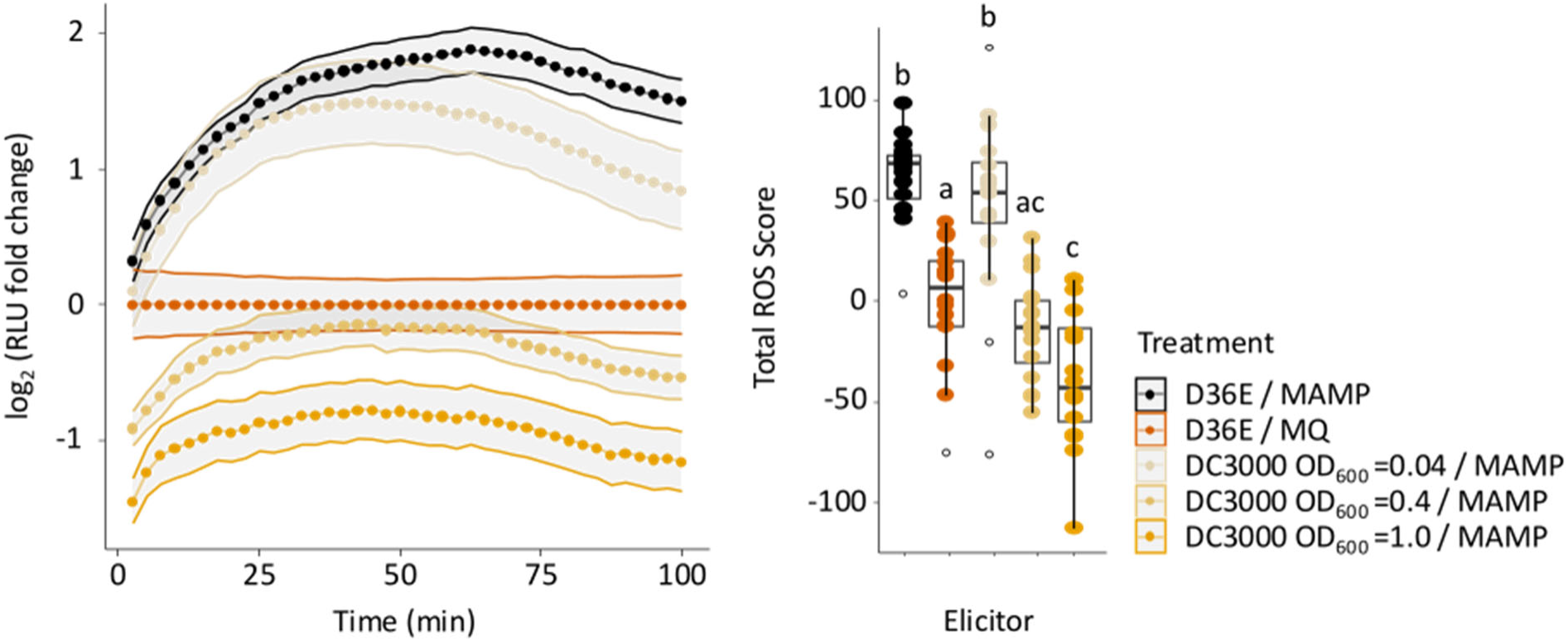
DC3000 supresses reactive oxygen species (ROS) bursts in spinach in a concentration-dependent manner. We infiltrated spinach with the ROS burst negative control D36E/MQ, the ROS burst positive control D36E/MAMP or DC3000 at different bacterial densities ranging from OD_600_ equal to 0.04, 0.4, to 1.0 plus the MAMP-mix. We show time-resolved graphs measuring ROS in relative light units (RLU) (**left**) and total ROS scores (**right**). Compared to the ROS burst negative control, D36E/MQ, DC3000 OD_600_ equal to 0.04 plus MAMP-mix showed only a trend, but no significant reduction of total ROS scores compared to D36E/MAMP, while DC3000 OD_600_ equal to 0.4 or equal to 1.0 plus MAMP-mix showed total ROS scores significantly lower than D36E/MAMP. Statistical analysis was conducted by ANOVA, followed by a Tukey’s Honestly Significant Difference test (*p* < 0.05). Letters above the boxplots indicate statistically significant different groups

**Figure S7:**
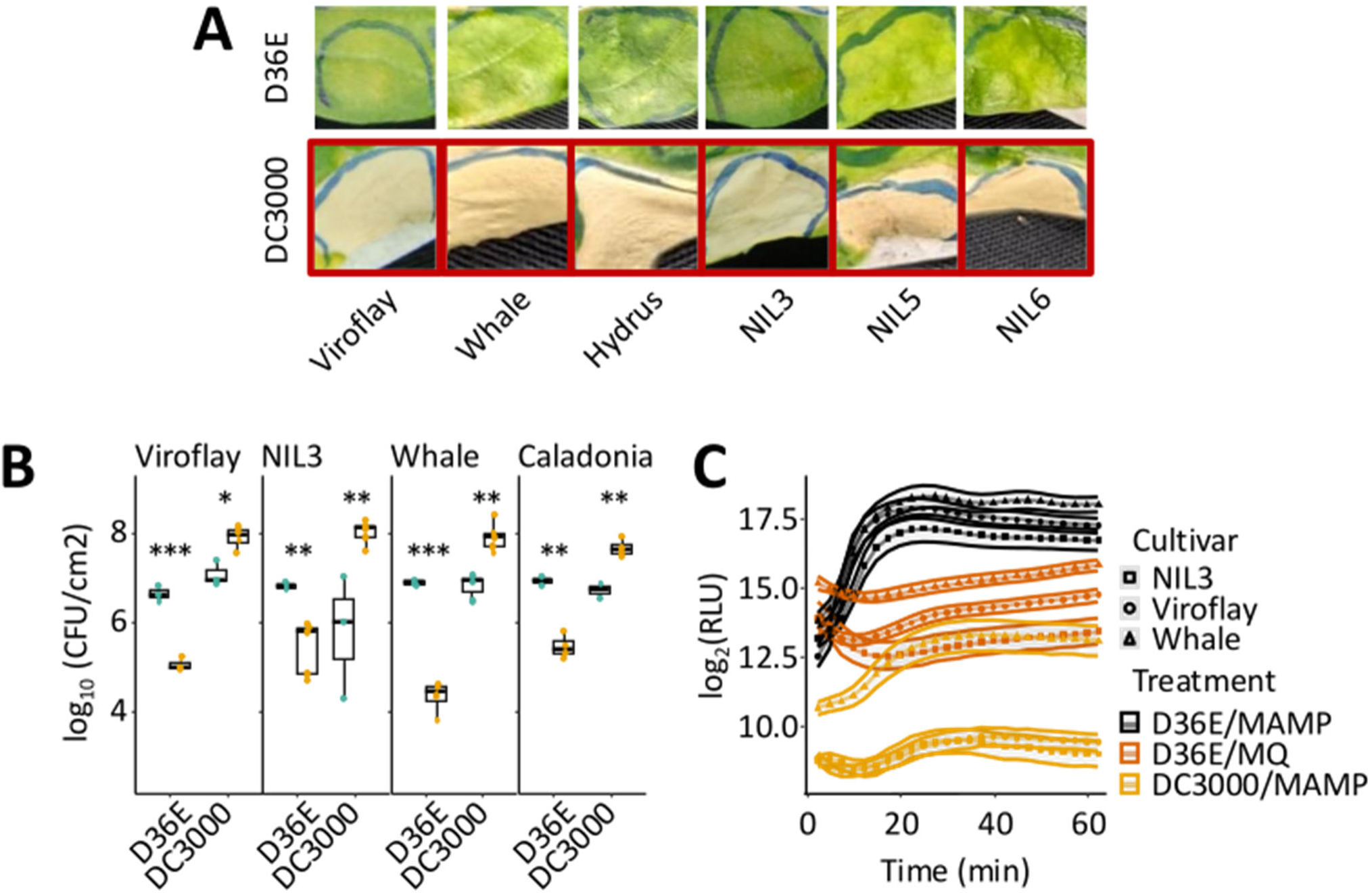
Different spinach cultivars react similarly to control treatments. We assessed if spinach cultivars differed in their response to control treatments with D36E or DC3000 in symptom development, bacterial proliferation or reactive oxygen species (ROS) bursts following MAMP elicitation. In **A)** we assessed symptom development in response to bacteria infiltration. All cultivars show a strong, cell-death/necrosis upon infiltration of DC3000 but not D36E. **B)** In the tested cultivars, D36E titres (colony-forming units or CFU) significantly decrease between 0-day post-infiltration (dpi) and 2 dpi, while DC3000 titres significantly increase over the same period. We applied a two-tailed t-test (*** p < 0.001, ** p < 0.01, * p < 0.05). **C)** One day prior to ROS elicitation with the MAMP mix, spinach was infiltrated with either D36E or DC3000 and ROS was measured in relative light units (RLU). In all three tested cultivars, D36E/MQ treatment resulted in stable or slightly increasing ROS levels over time. D36E/MAMPs treatment elicited a ROS burst followed by elevated ROS levels. In contrast, DC3000/MAMPs treatment reduced ROS levels relative to D36E/MQ, though absolute ROS levels varied among cultivars.

**Table S1.**
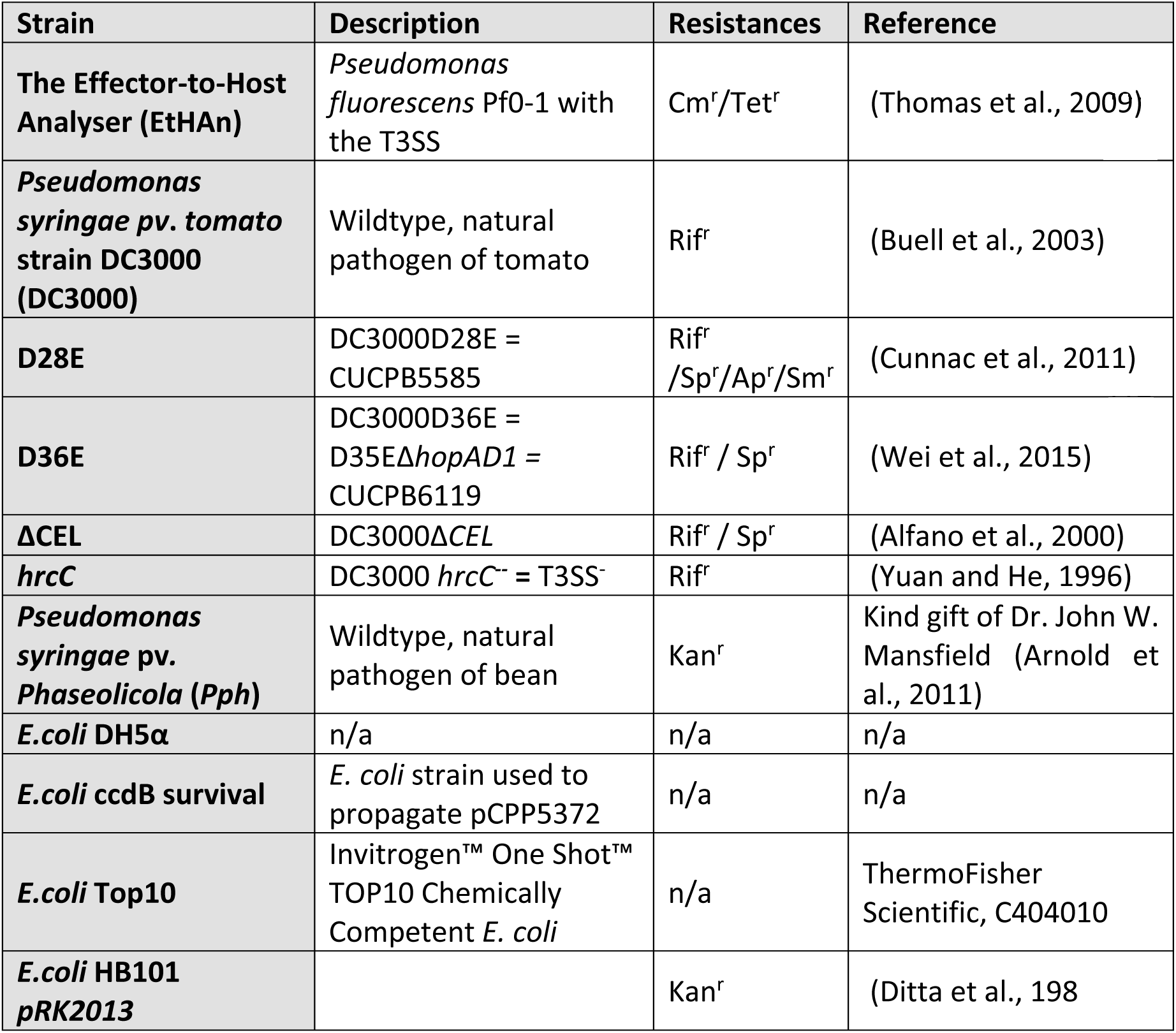
Detailed information on the bacterial strains used in this study.

**Table S2:** Antibiotics used in this study including abbreviations and standard working concentrations.

| Antibiotic | Abbreviation | Working concentration (µg/ml) |
| --- | --- | --- |
| Ampicillin | Ap | 100 |
| Carbenicillin | Cb | 100 |
| Chloramphenicol | Cm | 10 |
| Gentamycin | Gm | 25 |
| Kanamycin | Kan | 50 |
| Rifampicin | Rif | 50 |
| Spectinomycin | Sp | 50 |
| Streptomycin | Stp | 50 |
| Tetracycline | Tet | 20 |

**Table S3:** Plasmids utilized in this study. Including references and plasmid-encoded antibiotic resistances.

| Plasmid Name | Resistance | References |
| --- | --- | --- |
| pCPP5372 | GmR, CmR | (Oh et al., 2007) |
| DC300 Effector | pENTR constructs (Wei, 2018) | Resistance |
| AvrE1 | pCPP5912* | Kan<sup>r</sup> |
| AvrPto | pCPP5154 | Kan<sup>r</sup> |
| HopA1 | pCPP5530 | Kan<sup>r</sup> |
| HopB1 | pCPP5617 | Kan<sup>r</sup> |
| HopD1 | pCPP5588 | Kan<sup>r</sup> |
| HopE1 | pCPP5618 | Kan<sup>r</sup> |
| HopF2 | pCPP5058* | Kan<sup>r</sup> |
| HopG1 | pCPP5238 | Kan<sup>r</sup> |
| HopH1 | pCPP5306 | Kan<sup>r</sup> |
| HopI1 | pCPP5571 | Kan<sup>r</sup> |
| HopK1 | pCPP5099 | Kan<sup>r</sup> |
| HopM1 | pCPP5233* | Kan<sup>r</sup> |
| HopN1 | pCPP5060* | Kan<sup>r</sup> |
| HopO1-1 | pCPP5910* | Kan<sup>r</sup> |
| HopQ1-1 | pCPP3373 | Kan<sup>r</sup> |
| HopR1 | pCPP5951 | Kan<sup>r</sup> |
| HopT1-1 | pCPP3416 | Kan<sup>r</sup> |
| HopU1 | pCPP5328 | Kan<sup>r</sup> |
| HopV1 | pCPP5741* | Kan<sup>r</sup> |
| HopX1 | pCPP5534 | Kan<sup>r</sup> |
| HopY1 | pCPP3417 | Kan<sup>r</sup> |
| HopAA1-1 | pCPP5107 | Kan<sup>r</sup> |
| HopAA1-2 | pCPP5108 | Kan<sup>r</sup> |
| AvrPtoB | pCPP5616 | Kan<sup>r</sup> |
| HopAF1 | pCPP5539 | Kan<sup>r</sup> |
| HopAM1 | pCPP5052 | Kan<sup>r</sup> |
| HopAO1 | pCPP5537 | Kan<sup>r</sup> |
| HopAD1 | pCPP6450 | Kan<sup>r</sup> |

\* construct includes chaperone

**Table S4.**
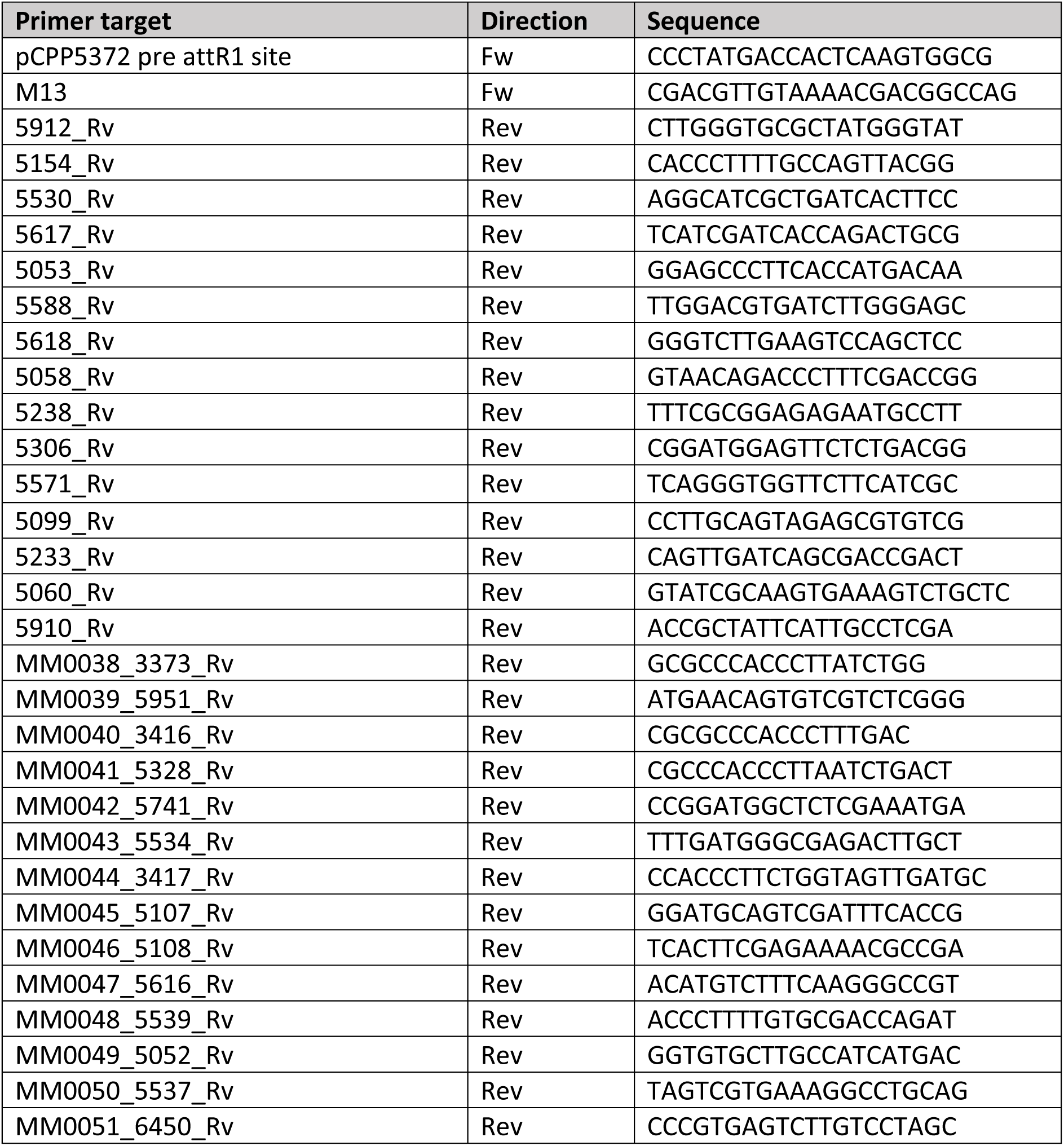
Primers utilized in this study. All primers were designed specifically for this project

**Table S5:** Elicitors tested for ROS burst elicitation on spinach.

| Elicitor-candidate | Description | Concen-tration | Producer/Source | Reference (if applicable) |
| --- | --- | --- | --- | --- |
| **Chitin** | Cell wall component of fungi, recognised by different receptors depending on the plant | 4 mg mL<sup>-1</sup> | Merck C7170, from shrimp shells, processing described in (Johannndrees et al., 2022) | (Sánchez-Vallet et al., 2015) |
| **flg22** | 22 amino acid peptide of *Pseudomonas aeruginosa* flagellin | 0.2 µM | Synthesised by Genscript; RP19986 | (Mersmann et al., 2010) |
| **nlp24** | 24 amino acid peptide of *Hyaloperonospora arabidopsidis* of the necrosis and ethylene-inducing peptide 1-like protein 3 (HaNLP3) | 2 µM | Synthesised by Genscript | (Oome et al., 2014) |
| ***P. effusa* 1 sporangia** | Heat-lysed *P. effusa* race 1 sporangia | 150 sporangia/µL | Sporangia harvested 14 dpi, cleaned with a vacuum filtration system combining Miracloth, spore collection on a 11 µm Merck Millipore Nylon net filter (NY1104700) | (Klein et al., 2020; Skiadas et al., 2022) |
| **MAMP-mix** | Heat-lysed D36E | OD<sub>600</sub> equal to 1.0 | Grown as described above, dilution/re-suspension in MQ | (Wei et al., 2015) |

